# Reorganization of motor functions within visuomotor networks subsequent to somatosensory cortical damage

**DOI:** 10.1101/2025.02.25.640230

**Authors:** Yuqi Liu, Elizabeth J. Halfen, Jeffrey M. Yau, Simon Fischer-Baum, Peter Kohler, Olufunsho Faseyitan, H. Branch Coslett, Jared Medina

## Abstract

Somatosensory inputs are critical to motor control. Animal studies have shown that primary somatosensory lesions cause sensorimotor deficits along with disrupted organization in primary motor cortex (M1). How does damage in primary somatosensory cortex (S1) influence motor networks in humans? Using fMRI, we examined two individuals with extensive damage to left somatosensory cortex, but primarily intact motor cortex and preserved motor abilities. Given left S1 damage, tactile detection and localization were impaired for the contralesional hand in both individuals. When moving the contralesional hand, LS, with near complete damage to the S1 hand area, showed increased activation in ipsilesional putamen and deactivation in contralesional cerebellum relative to age-matched controls. These findings demonstrate influences of S1 damage to subcortical sensorimotor areas that are distant from the lesion site, and a potential reweighting of the motor network with increased action selection in putamen and inhibition of sensory prediction in cerebellum in the face of sensory loss. In contrast, RF, who had a small island of spared S1 in the hand area, showed greater activation in contralesional S1 for movement versus rest. This same region was also activated by pure somatosensory stimulation in a second experiment, suggesting that the spared S1 area in RF still subserves sensorimotor processing. Finally, the right middle occipital gyrus was more strongly activated in both individuals compared with controls, suggesting a potential reliance on visual imagery in the face of degraded sensory feedback.

## Introduction

Successful hand actions critically depend on somatosensory feedback that informs the current state of the hand and the object. Integration of sensory information into motor commands relies on the communication between sensory and motor networks. The adjacent primary motor cortex (M1) and primary somatosensory cortex (S1) are densely interconnected and instantiate such communication (Porter & White, 1983; Ghosh & Porter, 1988; Mao et al., 2011; Tamè et al., 2015; Catani et al., 2012). Although simple tactile detection could be relearned (LaMotte & Mountcastle, 1979), lesions in S1 profoundly influence motor processing, reducing the power, dexterity and use of the contralateral hand in animals (Ghosh & Porter, 1988; Xerri et al., 1998; Mathis et al., 2017), humans (Carey et al., 2018; Chen et al., 2006; Jeannerod et al., 1984). Infarcts in S1 have altered response profiles (Kambi et al., 2011; Qi et al., 2010; Kato & Izumiyama, 2015) and organization in M1 (Harrison et al., 2013). These findings provide evidence for reorganization of primary motor cortex due to primary somatosensory damage.

Beyond primary motor cortex, motor networks encompass a wide range of cortical and subcortical areas. One dimension along which these areas vary functionally is the degree they participate in sensorimotor integration. For example, the cerebellum is thought to be critical for predicting sensory consequences during movement, and disrupting projections from S1 to cerebellum impairs motor performance in rats (Jenkinson & Glickstein, 2000). Recent studies established a double-dissociation between M1 and basal ganglia such that lesioning rats’ M1 affected the execution of visually-guided but not overly learned motor sequences, whereas damaging the basal ganglia impaired overtrained but not visually-guided motor sequences (Mizes et al., 2023a, 2023b). These findings suggest that M1 is required for integrating sensory information while the basal ganglia are essential for executing automatic, internally-generated kinematic patterns. One ramification of differential involvement in sensory processing is that each motor area may respond differently to damage in somatosensory cortex. However, the influence of somatosensory damage on motor processing in humans remains largely unknown.

To address this question, we examined two individuals with extensive damage to right somatosensory cortex, but largely intact motor cortex (**Figure 1**). Both individuals demonstrated impaired tactile detection and localization with relatively preserved motor abilities on the contralesional hand, providing a unique opportunity to investigate the reorganization of motor networks subsequent to somatosensory damage. Importantly, the two individuals differ in the extent of damage to the hand area of S1, with one individual (LS) having near complete damage and the other (RF) having a spared strip of S1 hand area along the central sulcus. These cases allow us to examine motor reorganization subsequent to somatosensory damage as well as the impact of the extent of the lesion.

**Figure 1.**
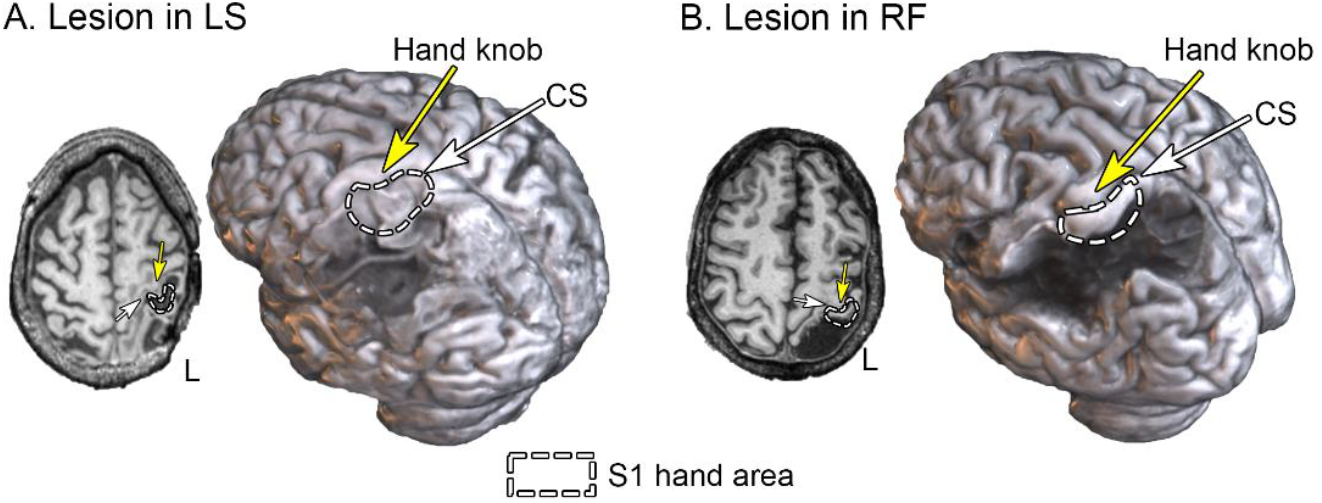
Lesion location in LS and RF shown in one transversal slice and a rendered 3D brain based on T1-weighted MRI scans. The yellow arrows point to the hand knob in M1, the white arrow points to central sulcus, and the dotted white outlines depict the estimated S1 hand area. Whereas LS had near complete damage to the hand area in S1, RF had an anterior portion spared. A detailed presentation of the lesions is shown in **Figure S1**.

We conducted an fMRI experiment in which LS and RF, along with eight age-matched controls, performed finger flexion-extension movements with each hand. Whole-brain comparisons between each brain-damaged individual and the control group were performed to examine changes in the activation pattern of motor regions as a result of somatosensory damage. In addition, to separate sensory processing from motor control, LS and RF participated in a second fMRI experiment where they passively received tactile stimulation on each hand. This experiment allowed us to examine whether the spared portion of S1 hand area in RF had remaining sensory function, and the influence of S1 damage on higher-level somatosensory areas (e.g., secondary somatosensory cortex) in LS.

## Results

### Impaired tactile abilities in LS and RF

Tactile detection was evaluated with Semmes-Weinstein monofilaments (North Coast Medical Inc., CA, USA) using a staircase procedure (see **Methods**). Both LS and RF showed elevated detection thresholds on the contralesional right hand (4g and 8g respectively), demonstrating diminished protective sensation. The ipsilesional left hand showed normal detection thresholds (0.07g for LS, 0.6g for RF; Hage et al., 1998).

In a tactile localization task in which the participants pointed to the tested hand where they felt touch, both LS and RF made significantly larger errors on the contralesional hand (LS: *localization judgment error* = 51.5 mm, *SD* = 29.0 mm; RF: *M* = 55.5 mm, *SD* = 32.6 mm) than on the ipsilesional hand ((LS: *M* = 10.5 mm, *SD* = 10.1 mm; RF: *M* = 5.3 mm, *SD* = 7.1 mm; permutation *p*s < .001; see **Methods** and **Figure S2A**).

An additional assessment of proprioception in LS demonstrated an impaired ability in localizing the contralesional hand when vision was unavailable (Contralesional hand: *localization judgment error* = 93.5mm, *SD* = 41.8mm, Ipsilesional hand: *M* =56.2mm, *SD* = 42.6mm, two-hand permutation *p* < .001; see **SI Methods** and **Figure S2B**). These findings suggest that the motor deficits on the contralesional hand of LS and RF when vision was not available likely stem from degraded somatosensory feedback. We then examined how the motor network in the brain responds to hand movement when somatosensory inputs are compromised.

### Contralesional motor activation

The motor fMRI experiment was a block design in which participants made periodic movements opening and closing their hand following visual cues (centrally presented “open” and “close”, **Methods**). The two hands were tested in separate runs. Motor activation was analyzed using standard general linear models (GLM) and indexed by the contrast of movement versus rest. LS, RF, and eight age-matched controls (all right-handed, mean age = 61.5 years, SD = 7.9 years, four females) participated in this experiment. Whole-brain voxel-wise Crawford-Howell t-tests were performed between each brain-damaged individual and the control group to examine quantitative differences. As data were collected from multiple sites, t-values of motor activation were used for this analysis to account for the internal noise associated with different scanners.

Brain damage induced by stroke is known to alter the shape of the hemodynamic response, at times causing a delay in perilesional brain regions (Amemiya et al., 2012; Bonakdarpour et al., 2007). Indeed, a lag analysis (**SI Methods**) showed that LS’s BOLD response to hand movement varied across brain regions (**Figure S3, S4**), ranging from no temporal shift (in putamen) to lagging by nearly six seconds (e.g., in motor cortex; **Figure 2A**). To account for the delay, we analyzed LS’s data in cortical regions using the FLOBS (FMRIB’s Linear Optimal Basis Sets) toolkit in FSL (West et al., 2019; **SI Methods**) with the initial delay of the HRF set to six seconds. As subcortical regions did not show such a delay, we modeled these regions using the standard HRF. No temporal delay was observed in RF (**Figure S3**).

**Figure 2.**
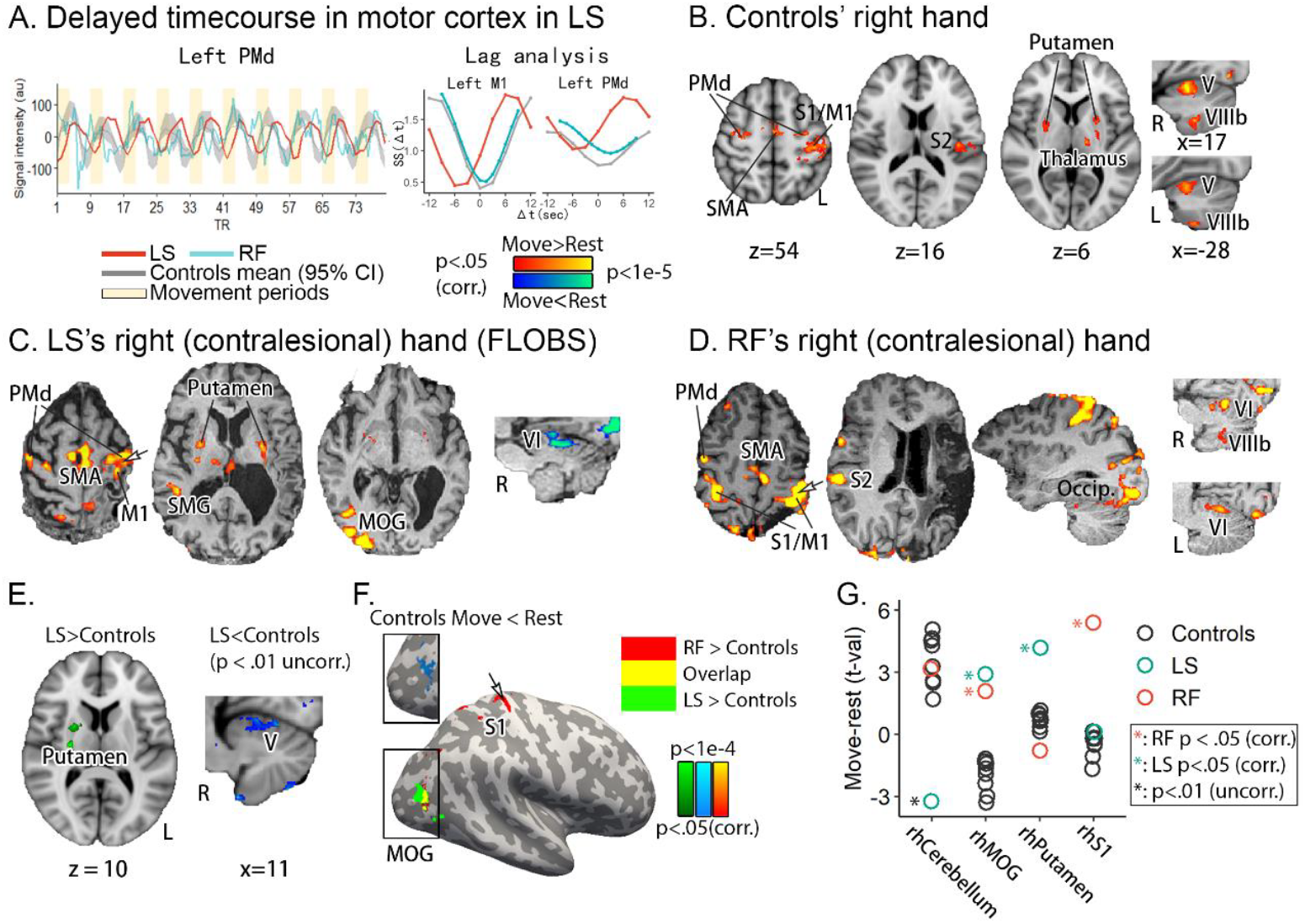
Brain activation when participants moved their right hand. A. Average (demeaned) time-course of the right-hand motor runs shows that LS’s hemodynamic response lagged behind controls’ and RF’s in PMd. Lag analysis revealed a delay of 6 seconds in LS’s hemodynamic response in M1 and PMd, as shown by the left-shifted red curve. B. In controls, moving the right hand significantly activated left primary somatosensory and motor cortex (SMC), bilateral PMd and SMA, bilateral putamen and cerebellum, left thalamus and S2. C. Contralesional motor activation in LS (FLOBS analysis). While showing significant activity in left M1, bilateral PMd, SMA, and putamen as with controls, LS showed significant activation in right MOG and deactivation in right cerebellum (standard analysis). D. In RF, other than left sensorimotor cortex, bilateral PMd and SMA, and cerebellum, RF showed significant activity in right S1 and M1, and right occipital lobe. Notably, the spared strip in S1 hand area was also activated. E. Whole-brain Crawford-Howell t-test. Based on the standard analysis, LS showed stronger activation in right putamen compared with controls. The right cerebellum was less activated than controls (ROI level p<.01 uncorrected). F. Both LS and RF showed significantly stronger activation in right MOG, an area that was significantly deactivated during movement in controls. RF also showed stronger activation in right S1. G: T values of each individual in areas found from the whole-brain Crawford-Howell t-tests. Asterisks reflect significance level from the whole-brain analysis. PMd: Dorsal premotor cortex. PMv: Ventral premotor cortex. SMA: Supplementary motor area. SMG: Supramarginal gyrus. MOG: Middle occipital gyrus. Occip.: Occipital lobe.

In controls, movement of the right hand activated cortical sensorimotor areas including contralateral left primary somatosensory and motor cortex (SMC), left S2, bilateral dorsal premotor cortex (PMd) and supplementary motor area (SMA; **Figure 2B**). Subcortical activation was found in bilateral putamen, left thalamus, and bilateral anterior (lobule V) and posterior (lobule VIIIb) cerebellum.

In LS, the same cortical motor areas – left M1, bilateral PMd and SMA, along with bilateral putamen – were activated as in controls (**Figure 2C**). No activation was seen in left S1 and S2 due to the lesion. Thalamic activation was also found, albeit on the contralesional right side. In addition, right supramarginal gyrus (SMG) and middle occipital gyrus (MOG) were activated, patterns not observed in controls. Finally, the standard analysis (with standard HRF) showed deactivation in lobule V in right cerebellum (see **Figure S5** for the time course and lag analysis). Whole-brain Crawford-Howell t-test between LS and controls revealed greater activity in right MOG (**Figure 2F**) and right putamen (**Figure 2E**). The cerebellar deactivation in LS manifested as significantly lower activity relative to controls (**Figure 2E**). The activity in MOG in LS is unlikely driven by visual stimuli (i.e. word instructions vs. rest) because controls showed less activity in this area during hand movement versus rest (**Figure 2F and 2G**) and the same trend of deactivation was seen during word presentation in a control experiment when no hand movements were made (*t*(6) = -2.42, *p* = .052; **SI Methods**).

In RF, moving the contralesional hand resulted in significant activity in bilateral PMd, SMA, as well as bilateral anterior cerebellum, as with controls (**Figure 2D**). There were three findings of note for RF. First, the spared contralesional S1, along with M1, was activated, consistent with controls. Second, ipsilesional sensorimotor cortex was also activated, whereas controls only showed contralateral activation. Finally, significant activity was found in right occipital lobe including MOG. In contrast to LS and controls, no significant activity was seen in bilateral putamen. Furthermore, ipsilateral right S2, as opposed to left S2 in controls, was activated. Whole-brain Crawford-Howell t-test between RF and controls revealed greater activation in right MOG, overlapping with where LS had stronger activity, and in contralesional right S1 (**Figure 2F**). The activation in MOG is located at the parietal-occipital sulcus and extends into Brodmann area (BA) 19 and BA37, close to and dorsal-posterior to extrastriate body area (EBA, **Figure S6**; Downing et al., 2001; Peelen & Downing, 2005).

To summarize, while both LS and RF showed greater activation in MOG relative to controls, LS demonstrated subcortical changes with higher activation in putamen and reduced activity in cerebellum, whereas RF presented cross-hemispheric changes with increased activity in contralesional S1 ipsilateral to the moving right hand.

### Somatosensory activation

The spared portion of left S1 in RF was activated by hand movement, potentially indicating preserved somatosensory functions in this area, yet tactile and proprioceptive abilities in RF were substantially impaired. Alternatively, the activation in S1 could be a spread of M1 activation due to spatial smoothing. To tease apart these possibilities, and to address the general question of how S1 damage influences the rest of the somatosensory network, we conducted a somatosensory fMRI experiment on LS and RF, with the ipsilesional hand serving as a within-subjects control. The experiment used a block design in which participants received periodic tactile stimulation on the palm of the tested hand (**Methods**). The contrast of touch versus rest was analyzed in each individual using standard GLMs.

Despite substantial damage in S1, both individuals reported feeling touch during the experiment. In RF, the spared portion of left S1, overlapping with motor activation, was also activated by tactile stimulation on the contralesional right hand (**Figure 3A**), suggesting residual somatosensory function in this area. In addition, several ipsilateral areas including PMd, PMv, and IPS were activated, overlapping with the left hand activation (**Figure 3B**). Tactile stimulation on the left hand activated a wide sensorimotor network including right sensorimotor cortex, right premotor and parietal cortex, bilateral LOTC, and bilateral cerebellum (**Figure 3B**).

**Figure 3.**
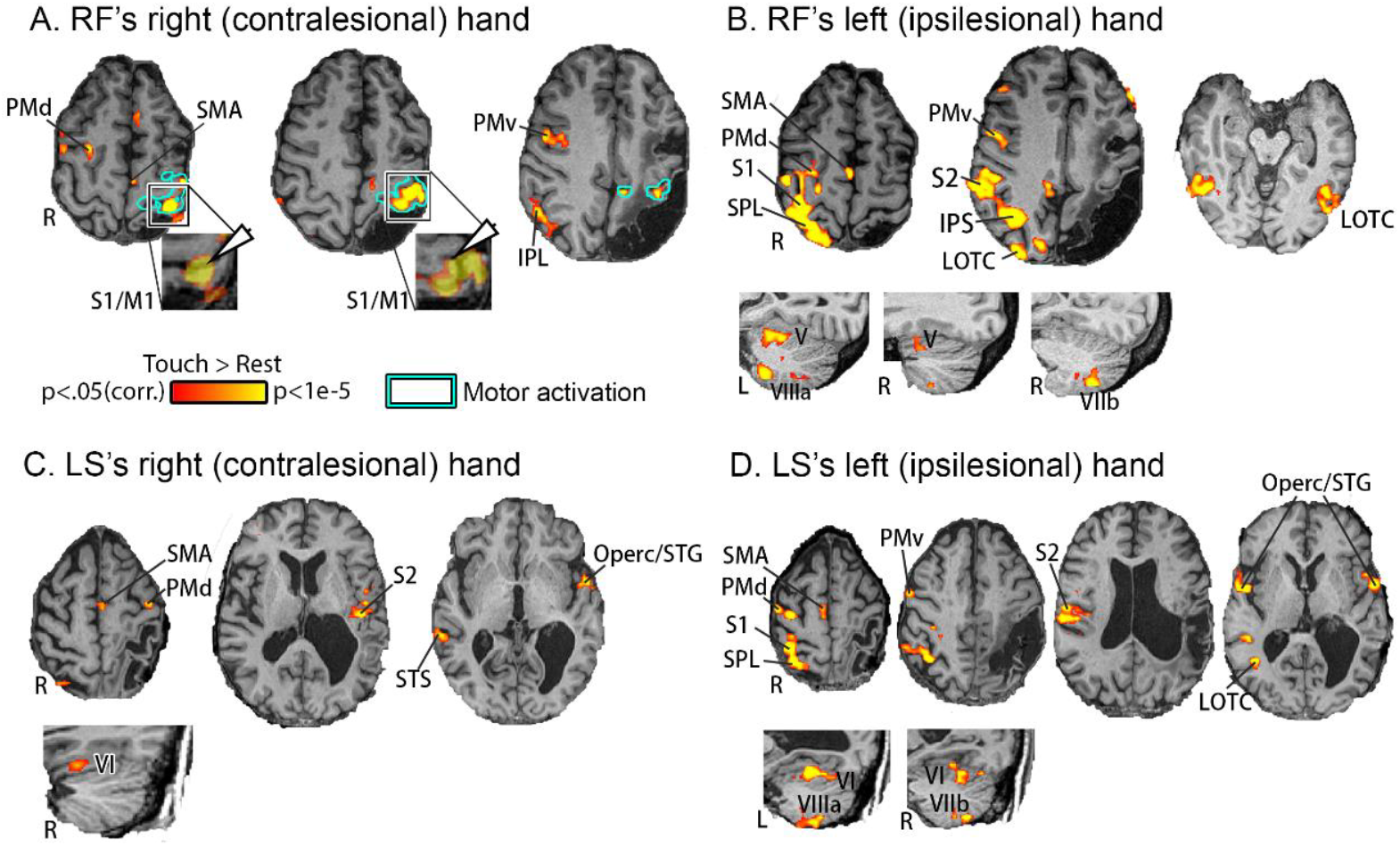
Somatosensory activation in RF and LS. A. Somatosensory stimulation on RF’s contralesional right hand activated the spared portion of left S1, overlapping with the motor activation. B. Tactile stimulation on RF’s left hand activated right S1, S2, PMd, PMv, SMA, SPL, IPS, and bilateral LOTC. C. Despite the lesion to S1, somatosensory stimulation on LS’s contralesional right hand activated higher-level sensorimotor areas including left SMA, PMd, S2, STG/frontal operculum, right STS and cerebellum, despite near-complete damage in S1. D. Similar to RF’s ipsilesional hand, tactile stimulation on LS’s left hand activated right S1, S2, PMd, PMv, SMA, SPL, bilateral STG/frontal operculum and cerebellum, largely consistent with LS. SPL: Superior parietal lobule. IPS: Intraparietal sulcus. Operc: Operculum. STG: Superior temporal gyrus. STS: Superior temporal sulcus. LOTC: Lateral occipital-temporal cortex.

In LS, no activation in left S1 was seen due to the lesion. Nevertheless, tactile stimulation on the contralesional right hand activated contralateral association areas including left S2, left PMd, left SMA, left frontal operculum, right superior temporal sulcus, and right cerebellum (**Figure 3C**), suggesting alternative pathways to S1. These areas are homotopic to areas activated by tactile stimulation on the ipsilesional left hand (**Figure 3D**).

### Ipsilesional hand movement

Finally, we present results when participants moved their left (ipsilesional) hand. In controls, moving the left hand activated right SMC, right PMd, bilateral SMA, and left S2 (**Figure 4A**). Subcortical activation was observed in right thalamus and bilateral cerebellum (**Figure 4A)**. The same cortical sensorimotor areas, namely right SMC, right PMd, and SMA, along with left cerebellum were also activated in LS (**Figure 4B**). LS also showed activation in ipsilateral left M1 and PMd, bilateral putamen, and bilateral MOG (**Figure 4B**). RF, with typical activation in right S1/M1, right PMd, and SMA, also demonstrated significant activation in bilateral occipital lobes that was not observed in controls (**Figure 4C**).

**Figure 4.**
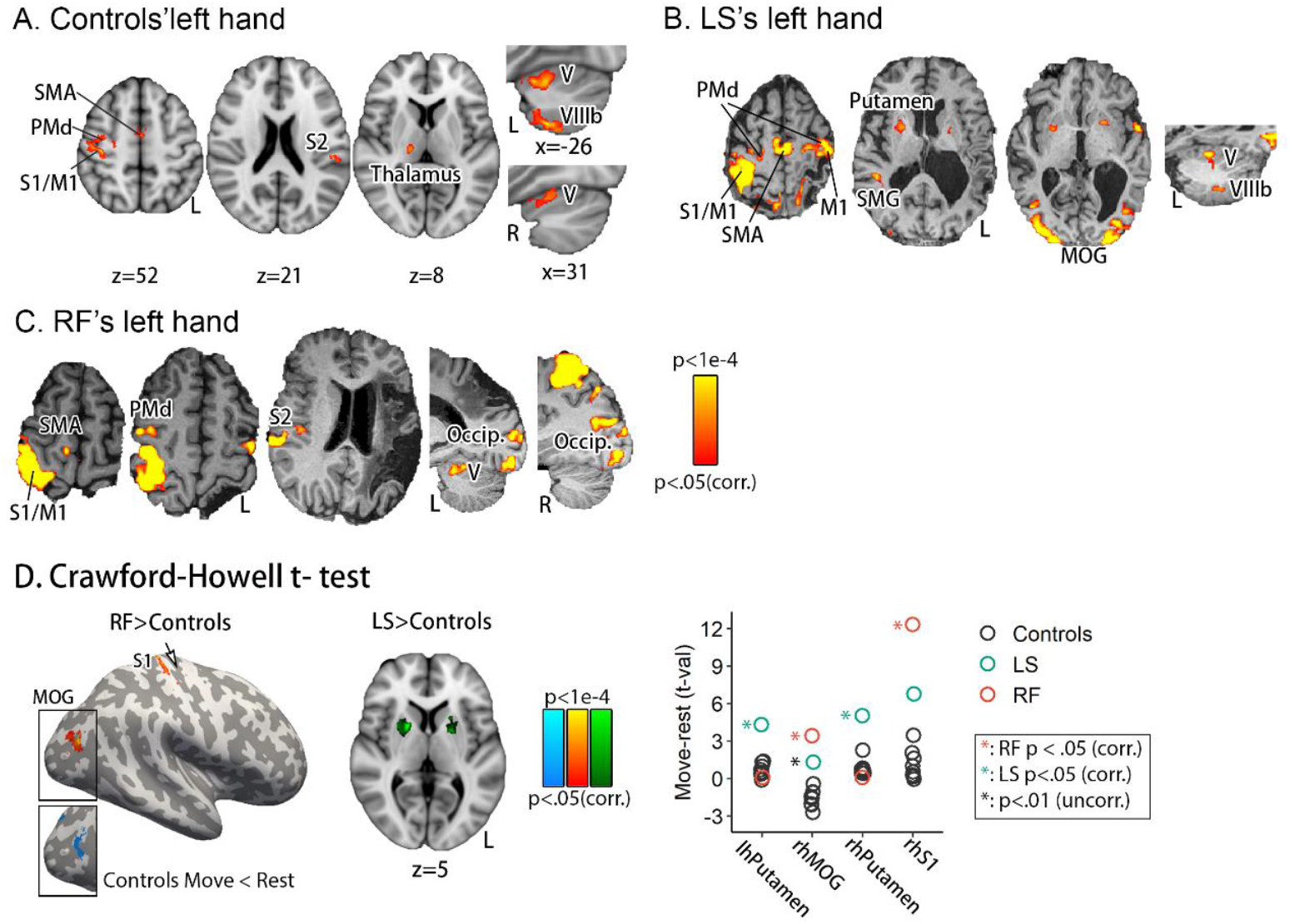
Brain activation when participants moved the left hand. A. In controls, moving the left hand activated right sensorimotor cortex (SMC), SMA, PMd, thalamus, left PMv, S2, and bilateral cerebellum. B. In LS, moving the ipsilesional left hand activated a similar motor network and additionally left and right MOG. C. In RF, moving the ipsilesional left hand activated typical motor networks and additionally right occipital lobe. D. Whole-brain Crawford-Howell t-tests revealed significantly greater activity in right MOG (where controls showed a significant deactivation) and S1 in RF relative to controls, and in bilateral putamen in LS relative to controls. ROI t-tests showed significantly stronger activity in LOTC also in LS relative to controls.

Crawford-Howell t-tests revealed similar patterns as observed for moving the contralesional right hand. Specifically, we found stronger activation in right S1 in RF (**Figure 4D**) and in bilateral putamen in LS (**Figure 4D**) compared with controls. Right MOG was more strongly activated in RF in the whole-brain analysis (**Figure 4D**), as well as in LS based on an ROI Crawford-Howell t-test (**Figure 4D**, point graph).

## Discussion

We examined two brain-damaged individuals to study the reorganization of somatosensory and motor functions after damage to S1. First, when moving their contralesional right hand, RF (small island of remaining S1) showed greater activation in ipsilateral right S1 compared with controls, while LS (no remaining S1 hand area) showed greater activation in right putamen and surprisingly, less activation in right cerebellum. Second, both LS and RF showed greater activity in right MOG during movement compared to controls. Third, tactile stimulation of the contralesional hand activated spared S1 in RF, whereas in LS activation was seen in higher-order sensory and motor areas.

### Cerebellar deactivation and increased activity in putamen in LS

Comparing LS with age-matched controls, we found greater activation in right putamen and less activation in right sensorimotor cerebellum, which was deactivated in LS during movement versus rest.

In right cerebellum, while controls showed significant activation during movement (lobules V-VI), LS demonstrated the opposite pattern – less activation during movement versus rest. During movement execution, the motor system predicts the sensory consequence of a motor command in compensation for delayed sensory feedback (Wolpert et al., 1995), a process proposed to be performed by the cerebellum (Nowak et al., 2007; Shadmehr & Krakauer, 2008; Blakemore et al., 2001; Izawa et al., 2012; Wong et al., 2019). The sensory prediction function involves both cerebellum and S1, regions that project to (Sultan et al., 2012; Glickstein, 1997) and modulate (Kilteni & Ehrsson, 2020; Matsui et al., 2012; Gold & Lauritzen, 2002; Restuccia et al., 2001) each other. Moreover, an absence of sensory input (due to neuropathies) but with intact S1 does not necessarily eliminate motor activity in cerebellum, suggesting the importance of S1 integrity in maintaining cerebellar activation (Weeks et al., 1999). In LS, we propose that deactivation of cerebellum during movement reflects inhibition of the sensory prediction function due to absence of sensory feedback and reduced input from S1.

An alternative interpretation is that the cerebellar findings in LS are due to hemodynamic lag due to the stroke (Amemiya et al., 2012; Bonakdarpour et al., 2007). We consider these results to more likely reflect deactivation for two reasons. First, hemodynamic lag is known to occur in perilesional but not remote areas. One study examined blood flow in patients with ischemia or hypoperfusion and found a delayed hemodynamic response in bilateral motor cortex but *not* in cerebellum (Amemiya et al., 2012). Second, although we found significant activation for somatosensory stimulation in the same cerebellar area for LS, no such lag was observed (see **Figure S3**). This suggests that our results are due to an actual change in cerebellar function in the face of somatosensory damage. Our results suggest a re-weighting of the motor system in compensation for absence of S1, wherein the relative weight was increased for the loop responsible for motor output (i.e., putamen), and decreased for the loop responsible for sensory processing (i.e., cerebellum).

The basal ganglia are active during action selection and decision-making, whereas cerebellum is responsible for estimating and processing the sensory consequences of movement (Doya, 2000; Jueptner & Weiller, 1998; Shadmehr & Krakauer, 2008). For example, the putamen is more strongly activated when performing self-timed versus externally-cued movements in non-human primates (Lee et al., 2006) and this activity is dramatically reduced for passive compared to active movements (Jueptner & Weiller, 1998; Liles 1985). Furthermore, patients with Parkinson’s disease, associated with basal ganglia degeneration, show substantial difficulty in performing voluntary actions versus visually-guided action (Jackson et al., 1995; see also Mizes et al., 2023). The increased putaminal activity in LS may reflect increased effort in selecting the correct motor plan (i.e., open or close the hand) under diminished somatosensory feedback.

### Increased activity in right S1 in RF

In RF, movement of the contralesional right hand activated typical motor networks. In addition, ipsilateral (to the moving hand) S1 was more strongly activated compared to controls, demonstrating reorganization to the contralesional hemisphere. Interestingly, such bilateral S1 activation was not observed in the somatosensory fMRI experiment (**Figure 3**), suggesting that it is specific to motor function. There is abundant evidence for the involvement of contralesional *motor* cortex when individuals with brain damage move the contralesional hand (Carey et al., 2006; Grefkes & Ward, 2014; Rehme et al., 2012; Ward et al., 2003; Favre et al., 2014), yet the involvement of contralesional *somatosensory* cortex remains unexplored. To our knowledge, this is the first report of contralateral somatosensory activity subsequent to S1 lesion. Increased activation in the contralesional hemisphere may result from decreased inhibition by the lesioned hemisphere or increased reliance on ipsilateral (to the moving hand) sensorimotor pathways (Grefkes & Ward, 2013; Medina & Rapp, 2008).

Why did LS and RF show substantially different activation patterns despite lesioned S1 and spared M1 in both cases? We speculate that these distinct reorganization patterns could be driven by different extents of their S1 lesions. Whereas LS suffered a complete lesion of the hand area in S1 and accordingly demonstrated no S1 activation for contralesional hand tactile stimulation, RF had a spared strip of S1 along the anterior bank that responded to touch. The spared tissue may represent residual tactile information from the hand, maintaining the connections between S1 and M1, basal ganglia, and cerebellum, thus preserving a typical motor network. Moreover, the spared S1 may have driven cross-hemispheric somatosensory reorganization via transcallosal connections (Grefkes & Ward, 2013). Animal studies found increased reorganization in remote areas (e.g., bilateral premotor cortex) with larger M1 lesions (Dancause et al., 2006; Dijkhuizen et al., 2003; Frost et al., 2003; Touvykine et al., 2016), and more substantial reorganization within S1 with more complete dorsal column lesions (Qi et al., 2019). We report for the first time from human subjects the effect of the extent of lesion size in the hand area in S1 on motor network organization. We note that the lesion extent also differs in other brain regions between LS and RF, with more severe damage in posterior parietal cortex (PPC) and inferior frontal gyrus (IFG) in RF compared to LS. While we cannot directly attribute the different motor activation patterns to specific lesions, the better manual abilities in RF, despite more severe lesions in PPC and IFG, indicates that the spared tissue in S1 might play a key role in the preservation of motor functions in RF.

### Increased MOG activity in both LS and RF

In both RF and LS, contralesional hand movement elicited activation in right MOG that was, surprisingly, significantly deactivated in controls. The MOG cluster is located dorsal-posterior to extrastriate body area (EBA), a region that responds to images of body parts versus other object categories (**Figure S6**; Downing et al., 2001; 2006; Weiner & Grill-Spector, 2011; Pilgramm et al., 2016). Although reported to be activated by the execution of hand and tool-use actions (Gallivan et al., 2011; Brandi et al., 2014; also see Hinkley et al., 2011), few studies have specified the function of MOG in motor control. We present two possible hypotheses for why it may be active in RF and LS. First, MOG has been associated with processing feedback (Bermann et al., 2012) and when the consequences of an action were incongruent with the participant’s intention (Yomogida et al., 2010). One possibility is that LS and RF relied more on active feedback monitoring relative to controls, thus activating MOG. Second, MOG has been associated with processing body stimuli, including viewing other people making gestures (Husain et al., 2009), inferring others’ action intention in a visual scene (Atique et al., 2011), and mental rotation of hand stimuli (Vingerhoets et al., 2002). A second possible explanation is that LS and RF adopt visual motor imagery in absence of sensory feedback (ter Horst et al., 2012; Mercier et al., 2008; Čeko et al., 2013), leading to stronger MOG activity. We note that the underlying mechanism of MOG involvement in motor control requires more experiments, yet our findings provide the first evidence that visual cortex compensates for sensory loss in motor control.

### Neural correlates of tactile perception after S1 damage

In LS, tactile stimulation on the contralesional right hand activated left S2 and frontal motor areas in absence of S1 activation. RF, however, showed activation in spared left S1 as well as right association areas. Both individuals reported feeling the touch during the experiment. Studies on both humans and non-human primates found that activity in higher-level areas such as SMA, PMd, and S2, but not in S1, coincides with conscious tactile detection (de Lafuente & Romo, 2005; Hernández et al., 2002; Grund et al., 2021; Moore et al., 2013; Tamè & Holmes, 2016). Along this line, sensory association areas may receive tactile inputs from different routes in LS and RF, giving rise to their tactile perception via different mechanisms. In LS, S2 may receive information from thalamic projections and then send this tactile signal to frontal motor areas (Friedman & Murray, 1986; Garraghty et al., 1991; Rowe et al., 1996; Hernández et al., 2002). In RF, the preserved S1 receives thalamic projections and feeds forward to higher-level areas. In either case, tactile detection is supported by existing neural substrates, but these substrates are insufficient to support accurate tactile localization.

In summary, we examined motor reorganization in two rare cases with substantially damaged S1 but primarily intact M1. LS, with a more profound lesion in S1 hand area, showed increased putaminal activation and cerebellar deactivation when moving the contralesional hand, suggesting re-balancing between motor output and sensory processing within motor networks. RF, with a strip of S1 hand area spared, demonstrated cross-hemispheric S1 reorganization. Both individuals showed increased activity in right MOG compared to controls, indicating compensatory visual imagery processes. These findings provide novel evidence for long-range motor reorganization and visual recruitment subsequent to S1 damage in human subjects, adding knowledge regarding the differential roles across motor regions in sensorimotor integration.

## Methods

### Participants

At the time of testing, LS was a 63-year-old male (seven years post-stroke) who suffered a left hemisphere infarct that extended from the central sulcus posteriorly to superior temporal sulcus (**Figure 1A**), affecting most of the lateral S1, part of S2, and inferior and superior parietal lobe. Upon observation, LS demonstrated clear deficits in touch and proprioception without vision, and heavily relied on visual information for moving the contralesional right hand. Precise manual tasks with vision (e.g., picking up a coin) were also difficult. He was clearly able to perform simple hand movements (e.g., opening and closing the fist) both with and without vision.

RF was a 49-year-old female (five years post-stroke) whose lesion, caused by an ischemic stroke, extended from the postcentral gyrus to superior temporal sulcus along with extensive damage to left parietal operculum, superior temporal gyrus, insula, inferior and middle frontal gyrus (**Figure 1B**). Critically, there was spared tissue in the anterior bank of the postcentral gyrus in RF, leaving the most anterior part of the hand area in S1 intact (**Figure 1B**). Although RF showed no noticeable manual motor deficit when vision was available, performing movements with the contralesional hand was difficult when vision was removed.

For the motor neuroimaging experiment, we tested eight age-matched neurologically typical control participants (all right-handed, mean age = 61.5 years, SD = 7.9 years, 4 females). This is a typical sample size in studies comparing single cases with control groups (e.g. four in Bonakdarpour et al., 2007; six in Weeks et al., 1999; seven in Amemiya et al., 2012, and 12 in Snow et al., 2015). All research was approved by the IRBs of Baylor College of Medicine (RF), University of Delaware (LS, controls) and University of Pennsylvania (LS), with all participants providing written informed consent.

### Behavioral testing

Tactile detection was examined on the palmar side of the tip of the middle finger, using a weighted 1-down 1-up staircase procedure (10 planned reversals for LS, 20 for RF) with Semmes-Weinstein monofilaments (North Coast Medical Inc., CA, USA, 20 monofilaments for LS ranging from 0.008g to 300g; five monofilaments for RF ranging from 0.07g to 300g). Starting from the heaviest filament, if the participant reported feeling the touch, the filament two levels lighter was used on the next trial, otherwise the filament one level heavier was used. Catch trials in which the experimenter approached the hand without touching were randomly interspersed in each block (approximately 1 out of 6 trials). The duration of tactile stimuli was approximately one second for tactile detection and localization. Tactile detection threshold was calculated as the mean filament intensity across all reversal points in the staircase procedure.

Tactile localization was assessed using clearly suprathreshold monofilaments using the method in Rapp et al., 2002 (300g on the contralesional hand of both LS and RF; 1g and 300g on the ipsilesional hand of LS and RF respectively, four filaments above the threshold in both cases). In each trial the experimenter touched one of 22 pre-determined locations on the hand dorsum (**Figure S2)** with the participant’s eyes closed. Then the patient opened their eyes and pointed with the other hand where they felt touch on the tested hand. A second experimenter coded their response on a standard hand template. Each hand was tested in two blocks with one trial per location in randomized order. Hand order was balanced in ABBA manner. Tactile localization error was calculated as the average straight-line distance (in mm) between the perceived location and the actual stimulus location. For each patient, localization performance on the contralesional hand was evaluated with the ipsilesional hand as a control by running a permutation test where the relationship between trial-wise localization judgments and hand was shuffled 100,000 times. The two-tailed permutation p-value was calculated as the percentage of permutations in which the absolute mean difference in localization error was larger than the actual absolute difference between the two hands. Permutation analyses were performed using DAAG package implemented in R (v4.1.1). Source data from behavioral tasks can be found at https://osf.io/zr6hb/.

### Neuroimaging experiments

#### Motor fMRI

Each run began with a baseline period of six seconds, then alternated between 12 seconds of movement and 12 seconds of rest for 10 cycles, totaling 246 seconds. In hand movement phases, the monitor, viewed from a mirror mounted on the head coil, flashed the word “open” and “close” at 0.5Hz and the participant opened and closed the tested hand accordingly. A central fixation cross was presented during rest. LS and controls ran three runs on each hand, RF ran two runs (due to time constraints) on each hand in an ABBA(AB) manner. Compliance with instructions was visually monitored during testing; all participants were easily able to complete the task.

#### Somatosensory fMRI

Each run began with a baseline period of six seconds, then alternated between 30 seconds of tactile stimulation and 30 seconds of rest for four cycles, totaling 246 seconds. During the experiment, the participant rested their hands on each side of the body, palms facing up. An experimenter manually stroked the palm with a brush at 2Hz frequency following a metronome presented via headphones that was only heard by the experimenter. The patient was instructed not to make any body movements and to fixate at a central cross on the monitor. Five runs were completed with LS (three contralesional, two ipsilesional due to time constraints), and four runs were completed with RF (two contralesional, two ipsilesional) in an ABBA(A) manner.

### MRI acquisition

LS’s neuroimaging data were acquired from a Siemens Tim Trio 3T scanner at the University of Pennsylvania. A structural image was collected using a T1-weighted MPRAGE sequence (TR=1620ms, TE=3.08ms, flip angle = 15°, 192*256*160 voxels, 1mm isotropic, FoV=19.2*25.6cm). Functional MRI images were collected using a T2^*^-weighted echoplanar imaging (EPI) sequence (TR=3000ms, TE=30ms, flip angle=90°, 64*64*48 voxels, 3mm isotropic).

Control participants were scanned using the same T1-weighted MPRAGE and T2^*^-weighted EPI sequences as with LS. Imaging data were obtained from a Siemens Prisma 3T scanner at the Center for Biomedical and Brain Imaging (CBBI) at the University of Delaware.

Imaging data from RF were obtained from a Siemens Prisma 3T scanner at the Center for Advanced MRI (CAMRI) at Baylor College of Medicine. A 3D anatomical scan was acquired using a T1-weighted MPRAGE sequence (TR=2300ms, TE=2.98ms, flip angle = 9°, 168*237*200 voxels (1mm isotropic)). Functional MRI volumes were collected using a T2*-weighted echoplanar imaging (EPI) sequence (TR=1500ms, TE=33ms, flip angle=90°, 96*96*69 voxels, 2mm isotropic).

### fMRI preprocessing

Neuroimaging data were preprocessed in FSL 6.0 (FMRIB’s Software Library). Preprocessing of functional scans involved removal of the first two volumes, high-pass filtering (cutoff at 100 seconds), slice-timing correction, spatial smoothing with 4mm full width at half maximum (FWHM), and motion correction through 6-degrees-of-freedom (DOF) rigid-body transformation to the middle volume. Head motion was within 1mm across all runs of all participants. T1-weighted structural images were brain-extracted with BET for controls and with optiBET (Lutkenhoff et al., 2014) for LS and RF. Brain-extracted T1-weighted images were then segmented into white matter, cerebrospinal fluid (CSF), and gray matter using FAST. Functional images were co-registered to brain-extracted structural images using 6-DOF transformation in FLIRT. Structural images were co-registered to the MNI152 template brain (1mm isotropic) using 12-DOF transformation in FLIRT for controls and using the LINDA package for LS and RF implemented in R (v4.1.1; Pustina et al., 2016). Finally, we reconstructed surface space for the MNI152 template brain in FreeSurfer (7.2.0) and projected results from volumetric to surface space for visualization.

### Statistical Analysis

#### Motor fMRI

Functional neuroimaging data were analyzed using a univariate general linear model (GLM). The blood-oxygen-level-dependent (BOLD) signal for each run was modeled with 16 regressors. The experiment regressor was created by convolving the box-car function modelling movement phases with a standard double-gamma HRF. This regressor implies the contrast of movement versus rest. The temporal derivative of the experiment regressor was added as a covariate of no interest to account for temporal variation in BOLD signal. LS’s data was additionally modeled with basis functions using the FLOBS toolkit to account for the temporal delay in motor cortex (**SI Methods**). An additional 14 nuisance regressors included six head motion parameters and their first derivative, mean time course from CSF, and mean time course from white-matter. Within each individual, runs of each hand were combined using a fixed-effects analysis. Group-analysis of controls was performed with a mixed-effects analysis using the FLAME 1 method in FSL. Results were thresholded at *p* < .001 and cluster-wise thresholded at *p* < .05 based on Gaussian Random Field (GRF) theory. Regions were identified based on atlases provided in FSL (Harvard-Oxford Cortical Structural Atlas, Juelich Histological Atlas, and Talairach Daemon Labels) and the Brodmann Area template provided in MRIcron.

To directly compare each brain-damaged individual with controls, we performed voxel-wise Crawford-Howell t-tests for each hand separately (Crawford & Howell, 1998) on the t-value of the movement-versus-rest contrast between each patient and controls across the whole brain, masking out the lesioned areas of each patient. For LS, this analysis was performed twice, once with results from the standard analysis and once with the FLOBS analysis for reasons discussed above. Results are thresholded at *p* < .01 and cluster-wise thresholded at *p* < .05 based on GRF theory. Individual *t*-values were extracted from the resulting regions and plotted to visualize the distribution. When noted, Crawford-Howell t-tests were performed at the ROI level to explore effects that may not survive whole-brain multiple comparisons correction.

#### Somatosensory fMRI

Data were analyzed using the same pipeline as with the motor experiment, except the experiment regressor was generated based on the design of the somatosensory experiment.

## Acknowledgements

We are grateful to LS and RF and all control participants for their dedication to this project. We would like to thank Emily Baumert, Michael Grzenda, Alexandria O’Neal, and Charlotte Wilkinson for their work on this project. This work was supported by grants from the National Natural Science Foundation of China (32200843 to Y.L.) and the National Science Foundation (1632849 to J.M.). This work is partially based upon work conducted by S. F-B. supported by the Independent/Research Development program while serving at the National Science Foundation.

