## Supplementary material for "Reorganization of motor functions within visuomotor networks subsequent to somatosensory cortical damage": SI Methods, Figure S1, Figure S2, Figure S3, Figure S4, Figure S5, Figure S6

**Supplementary Information**

**Methods**

**Proprioceptive hand localization in LS**

LS’s tested hand was placed on a slider on a table. The hand and slider were covered under a white board. We instructed LS to rest the tested hand on the slider without making any voluntary movements. In each trial, the experimenter moved the slider to align LS’s middle finger with one of the 15 locations (arranged in a grid of 30 cm wide and 16 cm deep, aligned with the body midline) under the white board. Then LS pointed to a location on the white board directly above the perceived middle fingertip location using the untested hand. A picture of each judgment was taken and later compared with the actual target location. Each location was tested once in each block. Each hand was tested in two blocks in an R-L-L-R order. Permutation tests were performed between the two hands on the straight-line distance (in mm) between the perceived and the actual hand location (**Main** **Methods**).

**fMRI visual control experiment**

During the motor fMRI experiment, participants viewed the words “open” and “close” during hand movement versus a fixation cross during rest, raising the possibility that motor activation were driven by viewing words. To examine the effect of word presentation alone, control participants performed two runs of a visual control experiment in which they passively viewed the same visual presentation as in the motor experiment (i.e. 12-seconds of word presentation, 12-seconds of fixation cross) while no hand movements were made. The contrast of word versus fixation cross was computed within each control participant and a one-sample t-test was conducted on the t-values within the MOG area where both RF and LS showed greater motor activity than controls (**Figure 2F**, overlap). Seven out of the eight controls participated in this experiment. One control participant did not complete this experiment due to time constraints.

**Assessing BOLD time course in LS and RF**

Brain damage induced by stroke is known to alter the shape of the hemodynamic response, at times causing a delay in perilesional brain regions (Amemiya et al., 2012; Bonakdarpour et al., 2007). In this case, the standard hemodynamic response function (HRF) would not accurately model the brain response. To assess the time course of the HRF in LS and RF, we performed a whole-brain lag analysis in each individual under each task as in Amemiya et al., 2012. For each voxel, the normalized root-mean-square (RMS) between its time course shifted at each Δt and the reference time course was calculated, with the reference time course derived by convolving the standard HRF with the box-car function corresponding to the experimental design. The Δt at which the normalized RMS reaches the minimum was deemed the temporal shift of the voxel’s time course relative to the reference.

**FLOBS analysis**

As it is unlikely that hemodynamic lag in perilesional regions would affect subcortical or cerebellar areas, we modelled BOLD signal using the standard HRF in these regions in LS. However, as shown in the whole-brain lag analysis, LS’s BOLD response to hand movement varied across brain regions (**Figure S3**), ranging from no temporal shift (in putamen) to lagging by nearly six seconds or more (e.g., motor cortex). The six-second lag had a dramatic impact such that no activation in motor cortex was found when modeled with the standard HRF (**Figure S4**), yet there was a clear motor-evoked BOLD response (**Figure 2A**, **Figure S4**). To account for the delay, we additionally analyzed LS’s motor data in cortical regions using the FLOBS (FMRIB’s Linear Optimal Basis Sets) toolkit in FSL (West et al., 2019). This method allowed us to specify the range of parameters that determine the shape of the HRF. A set of basis functions was generated whose weighted combinations span possible HRFs that satisfy the specified parameters. We set the initial delay of the HRF to range from 0 to 6s to account for the delay in LS’s BOLD signal and selected the first six basis functions that collectively explain more than 95% of the variance of the sample HRFs. Each run of LS’s data was modeled with these six basis functions along with nuisance regressors. Significance of the first basis function regressor, which has the shape of a canonical HRF and implies motor activity versus rest, denotes motor activation level. To assess model performance, we sampled key regions of interest and calculated percentage variance explained (R-square) of the experiment regressor and of the full model of the FLOBS analysis and compared with the standard analysis. This assessment verified improved model fit by FLOBS compared with the standard analysis in LS’s motor cortex that was specifically contributed by the experiment regressor (**Figure S4).**

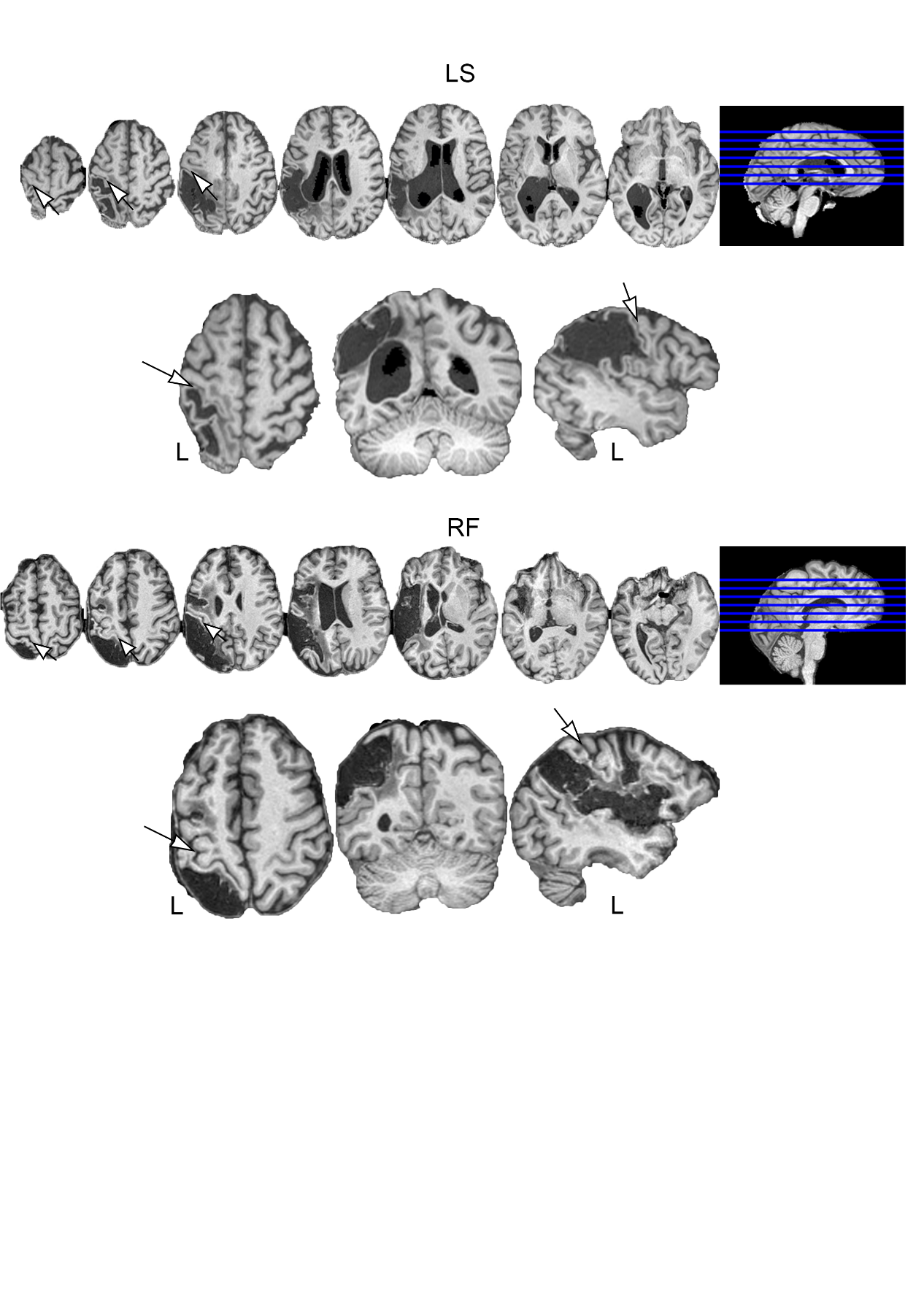

**Figure S1**. T1-weighted MRI scans of LS and RF. LS’s lesion extends from central sulcus over the posterior parietal cortex, sparing the motor cortex. RF’s lesion extends from postcentral cortex to posterior parietal cortex along with damage in superior temporal cortex, inferior frontal gyrus, and middle frontal gyrus. The motor cortex and the anterior strip of S1 is spared.

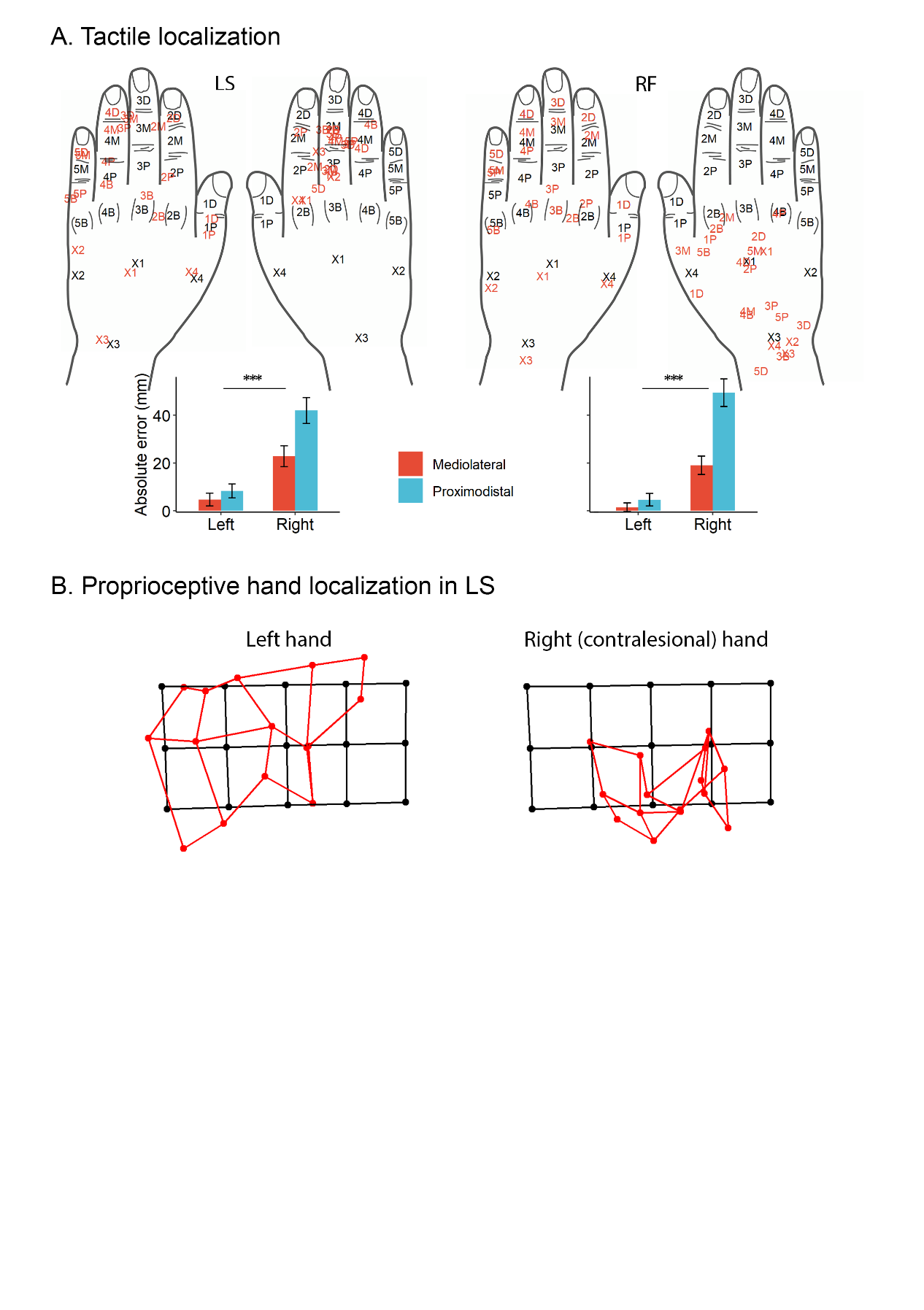

**Figure S2**. A. Tactile localization performance of LS and RF. Black labels mark target location, and red labels mark perceived location. Both participants made significantly larger errors on the contralesional right hand compared to the ipsilesional left hand. B. Mean arm localization performance in LS. Black dots are actual locations of the middle finger, red dots are perceived locations. LS made larger localization errors on the contralesional right hand than the ipsilesional left hand (permutation *p* < .001).

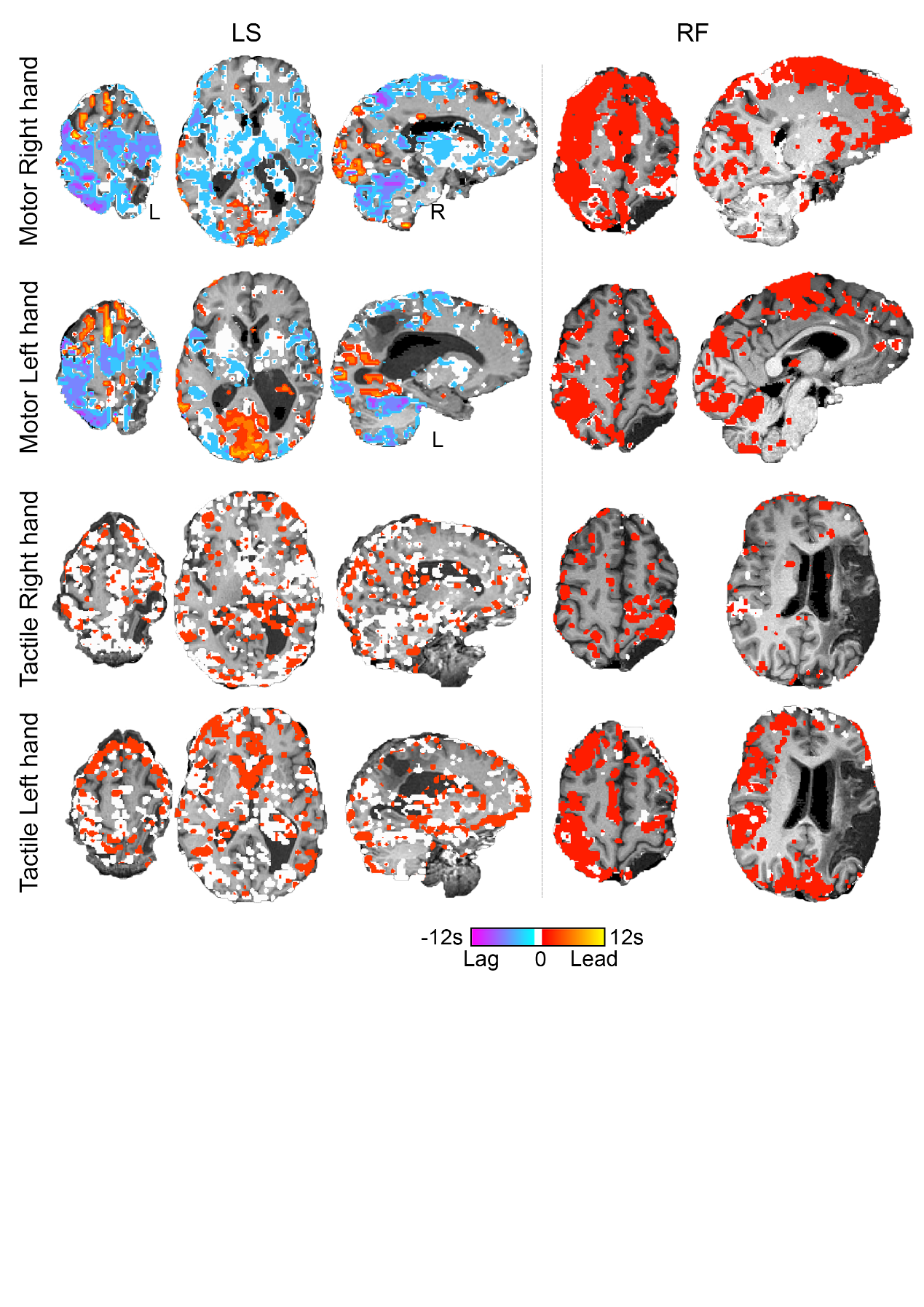

**Figure S3**. Whole-brain lag analysis between each voxel’s time-course and the reference time-course obtained by convolving the experimental conditions with the standard double-gamma function. Voxels that are i) within the lesion and ii) did not show significant correlation with the reference at all temporal shifts were masked out. During the motor task, both hands of LS showed lagged hemodynamic response in motor cortex and cerebellum, but not in bilateral putamen. No substantial temporal shift was found in LS’s tactile response, nor in RF’s tactile or motor response.

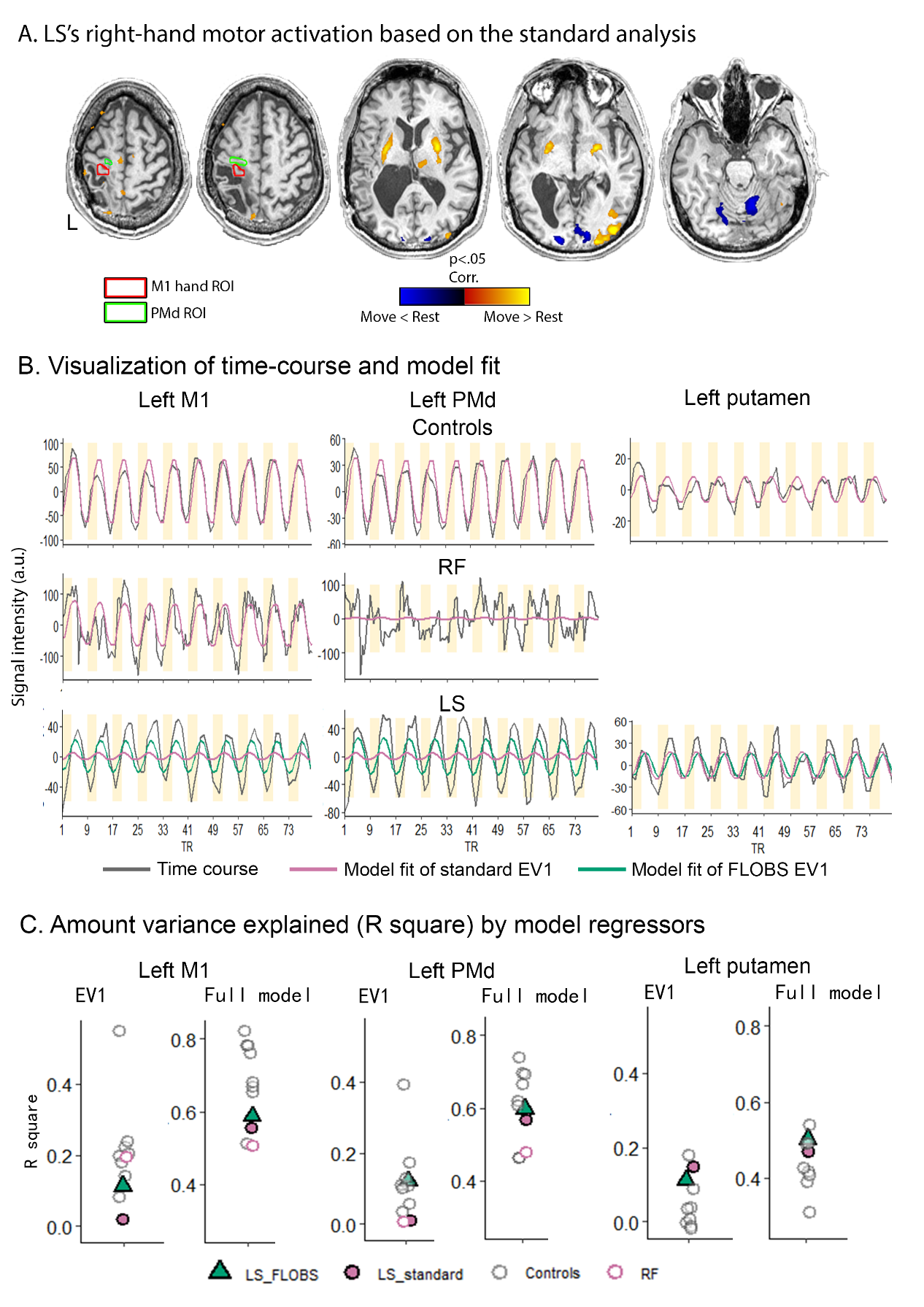

**Figure S4**. A. Motor activation of LS’s right hand mapped using the standard HRF. No activation was found in motor cortex due to the temporal shift in LS’s hemodynamic response. M1 hand ROI (outlined in red) and PMd ROI (outlined in green) are areas showing significant activation using the FLOBS analysis (see B.). B. Time-course and model fit in left M1 hand area, left PMd, and left putamen in LS, RF, and controls. For left M1 and left PMd, the first EV of FLOBS fits LS’s time-course better than the standard EV. Left putamen in RF was not plotted because RF did not show activation there, hence there was no event-locked time-course to inspect. C. Amount of variance explained (R-square) by different regressors in each ROI. For LS, the first EV of FLOBS improved modelled fit (i.e., increased R-square) relative to the standard EV in left M1 and left PMd (green triangle vs. magenta circle under the EV1 plot). The improvement is specific to the experimental EV because the R-square of the full model remains comparable. On the other hand, left putamen, showing no temporal delay, is better explained by the standard EV.

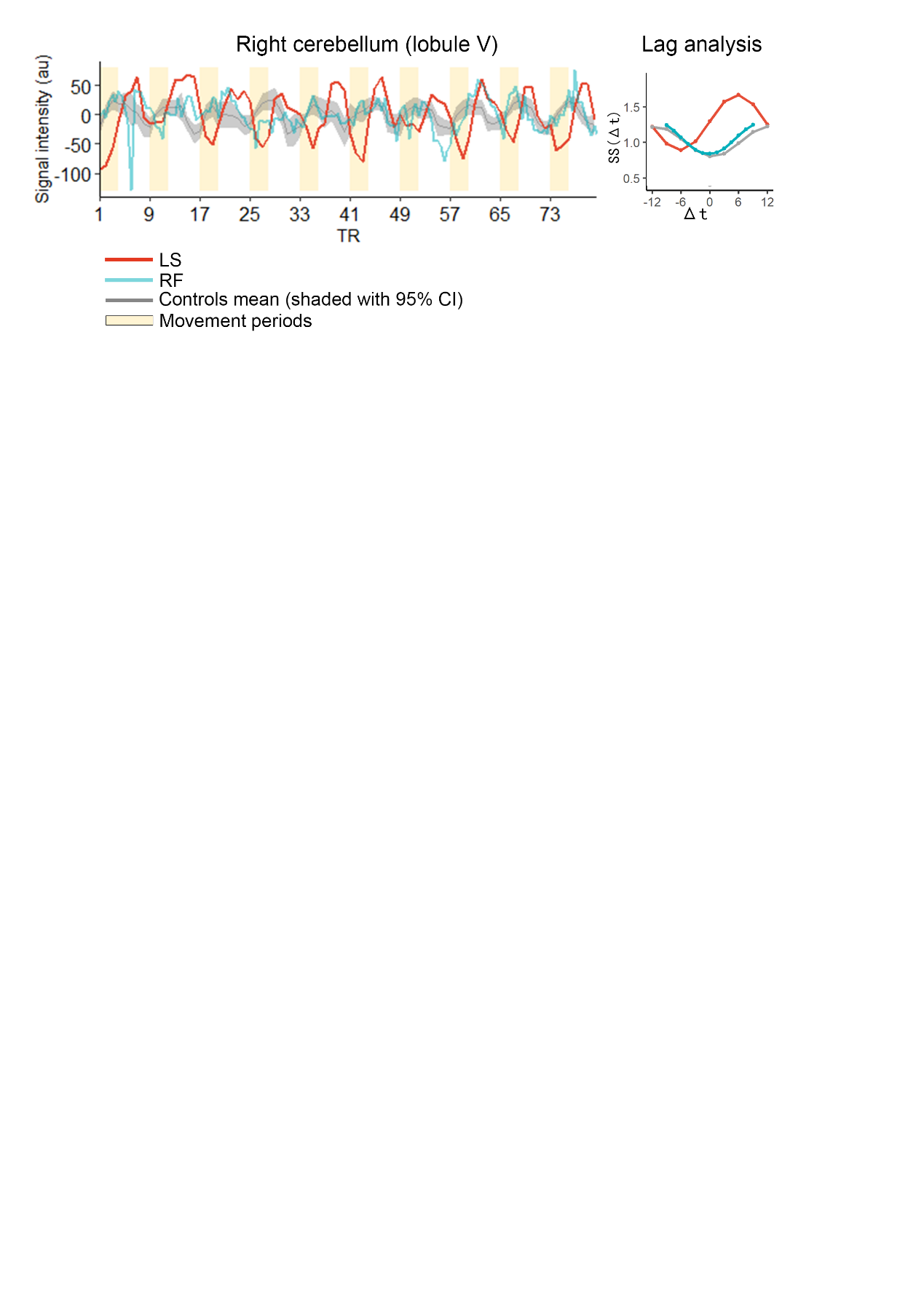

**Figure S5.** Time course in right cerebellum (lobule V) of LS shows a delay relative to movement onset and relative to RF and controls. Lag analysis confirms the delay in LS but not in RF or controls.

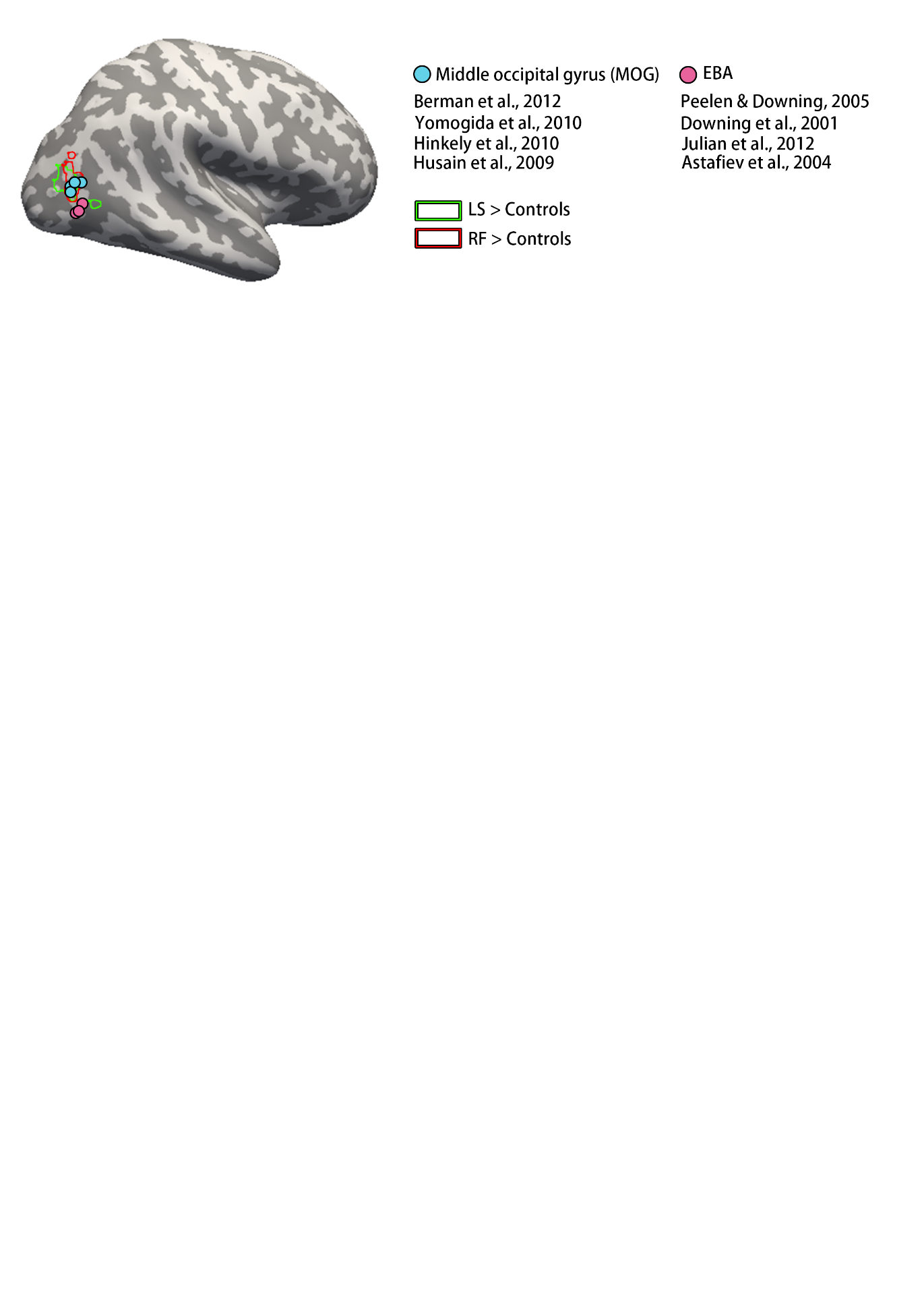

**Figure S6.** The visual area where both LS and RF showed greater activation when moving the contralesional (right) hand compared to controls is dorsal-posterior to extra-striate body area (EBA) and corresponds to middle occipital gyrus. The order of the references matches the ventral-dorsal arrangement of the dots on the brain surface.
